# Reconstruction of distinct vertebrate gastrulation modes via modulation of key cell behaviours in the chick embryo

**DOI:** 10.1101/2021.10.03.462938

**Authors:** Manli Chuai, Guillermo Serrano Nájera, Mattia Serra, L. Mahadevan, Cornelis J. Weijer

## Abstract

The morphology of gastrulation driving the internalisation of the mesoderm and endoderm differs dramatically among vertebrate species. It ranges from involution of epithelial sheets of cells through a circular blastopore in amphibians to ingression of mesenchymal cells through a primitive streak in amniotes. By targeting signalling pathways controlling critical cell behaviours in the chick embryo, we generated crescent- and ring-shaped mesendoderm territories in which cells can or cannot ingress. These alterations subvert the formation of the chick primitive streak into the gastrulation modes seen in amphibians, reptiles and teleost fish. Our experimental manipulations are supported by a theoretical framework linking cellular behaviors to self-organized multi-cellular flows in the accompanying paper. All together, this suggests that the evolution of gastrulation movements are largely determined by the shape of and cell behaviours in the mesendoderm territory across different species, and controlled by a relatively small number of signalling pathways.

---

During gastrulation, embryos transform from a simple epithelial layer into a three-layer structure. In vertebrate embryos, this process requires large morphogenetic movements that locate the three primary germ layers, the ectoderm, mesoderm and endoderm, in their topologically correct positions (1). However, the detailed tissue movements and morphology of the embryo during gastrulation vary greatly among vertebrate animals (1). Frogs form a blastopore, through which the mesoderm and endoderm precursors internalise as epithelial sheets while, at the other end of the spectrum, chick and mouse embryos internalise these precursors as mesenchymal cells through a structure known as the primitive streak (2).

The morphogenetic tissue flows in all organisms result from the integration of critical cell behaviours including cell division, differentiation, contractility and movement. These cell behaviours are patterned and coordinated through short and long-range chemical and mechanical signals (3, 4). Different spatiotemporal organisation of these cell behaviours, in conjunction with geometrical and mechanical constraints imposed by the size and shape of the embryo and extraembryonic tissues, determine the diverse morphogenetic programs in vertebrate gastrulation. However, how the large scale cell movements are coordinated and coupled via feedback to the signalling processes as part of robust developmental control mechanisms remain to be resolved (5).

Using the chick embryo as a model organism, we explore how the patterning of cell behaviours determine primitive streak morphogenesis during gastrulation (Fig. 1A-C, G). To study the tissue-wide coordination of the cell behaviours in the epiblast, we image gastrulation in a chick line expressing a membrane-targeted GFP using a custom build light sheet microscope (6) and image analysis (movie 1; red titles are hyperlinks to the movies. Click to open the movies in the browser). The dynamics of tissue deformation (Fig. 1D) can be directly quantified by measuring changes in the velocity field associated with tissue shape changes and movement using particle image velocimetry (PIV, Fig. S1A-B). The decomposed strain rates calculated from the velocity fields are a robust indicator of individual cell behaviours (Fig.1E, Fig. S1C, see methods). Negative values in the isotropic part of the strain rate (associated with convergence) denote regions where cells are undergoing apical contraction and cell ingression, and areas characterised by large values in the anisotropic components of the strain rate (associated with shear) indicate the convergent extension of the tissue produced by directed cell intercalations. Our observations show that the primitive streak forms in a region undergoing convergent extension produced by apical contraction, cell ingression and cell intercalation (Fig. 1E).

**Fig. 1:**
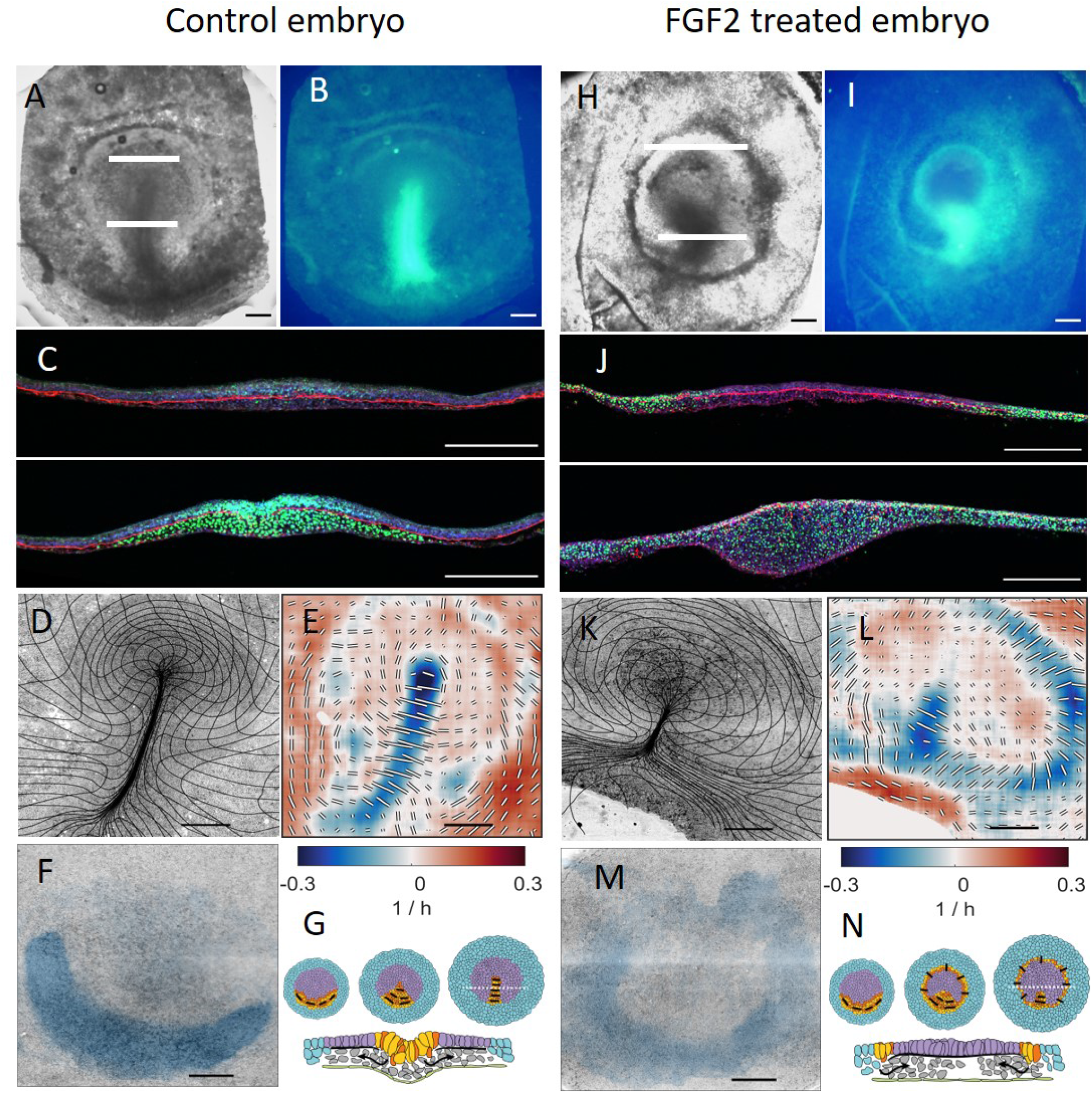
Characterisation of gastrulation for control embryo (A-F) and after FGF2 treatment (H-N). A: brightfield image of embryo at HH3+. B: Slug expression in embryo shown in A. C: Two sections taken at white lines in A showing slug (green), fibronectin (red) and actin (blue). Note that in the streak the fibronectin layer is fragmented. D: Overview images of embryo from lightsheet microscope at the end (HH3+) of the experiment overlaid with a deformation mesh. E: Strain rate tensor for stages shown in D. isotropic strain rate component shown as colour, blue for contraction, red for expansion. The anisotropic part of the strain rate tensor is shown as black and white bars indicating magnitude and direction of contraction. F: Cells that will ingress through the streak (domain of attraction) during the experiment calculated from the dynamic morphoskeleton. G: Schematic summary of development (stages HH1, HH2, HH3), extraembryonic area blue, mesendoderm precursors yellow/red embryonic area purple, black bars show direction of cell intercalation, section taken at broken white line in HH3 embryo schematic E. Scale bars 500 *µm* in all images except in C, where the scale bar is 250 *µm*. For panels E and L the scale bar length represents ∼ 0.5 1/h for the anisotropic component of the strain rate. Panels H-N same layout as panels A-F for FGF2 treated embryo for the same time as the control embryo.

To quantify the scale and nature of these movements requires an effective description of the relative motions of cells. While Eulerian quantities (decomposed strain rates) identify distinct cell behaviors, morphogenetic features arise from different processes integrated over space and time along cell paths. To account for this, we compute the dynamic morphoskeleton (DS) (7), which consists of Lagrangian attractors, their domain of attraction (DOA) and repellers that altogether organize tissue flows (Fig. 2 and movie 3 of the accompanying paper (8)). The DOA consists of the back projection of the attractor on the initial stage of development and reveals the extent of the sickle-shaped posterior located mesendoderm precursors fated to ingress through the streak (Fig. 1F). See methods for details on DM computation.

**Fig. 2:**
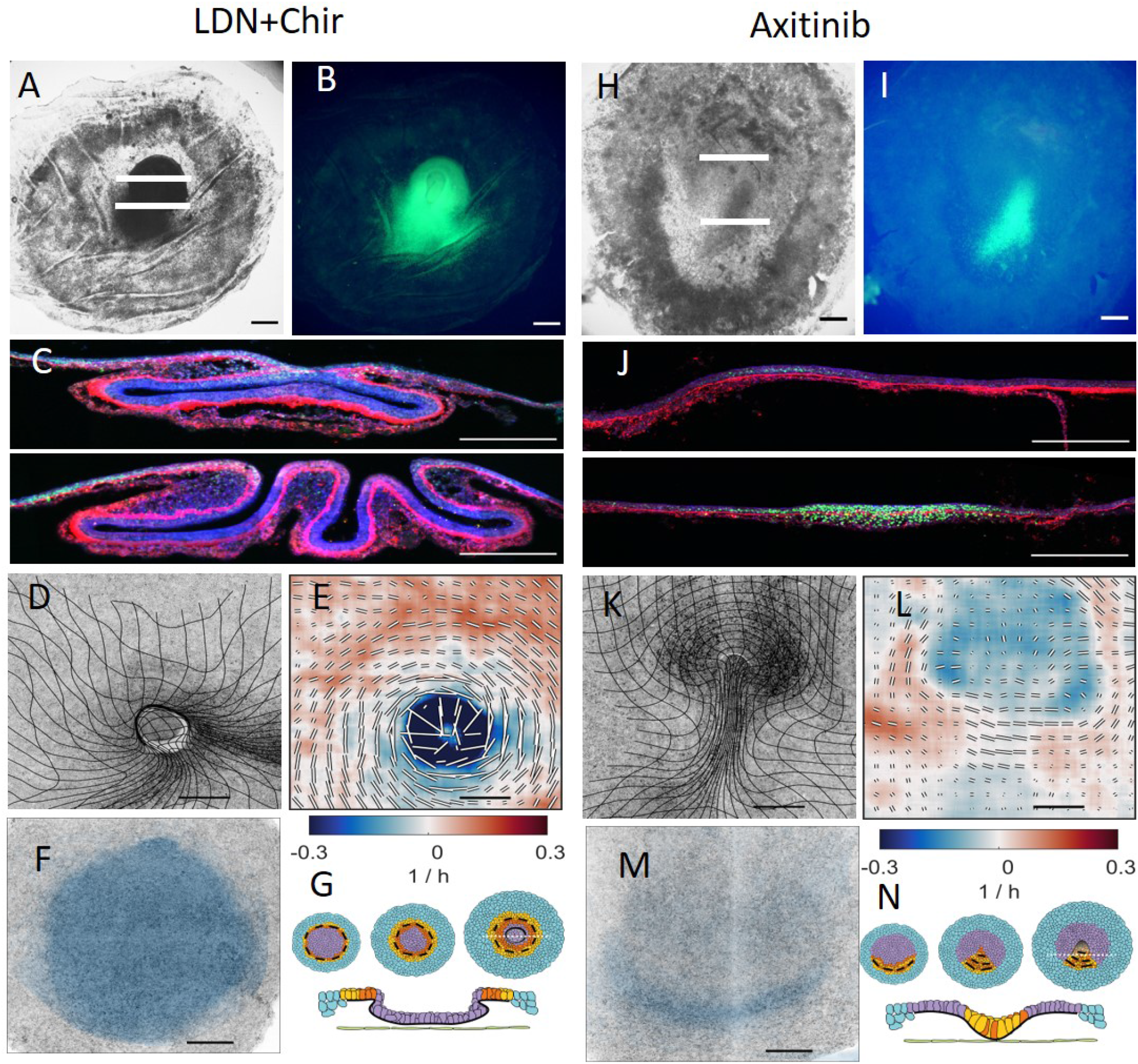
Characterisation of gastrulation for embryo treated with LDN-193189 and CHIR (A-F) and in an embryo treated with Axitinib (H-N). Embryos were treated with the BMP receptor inhibitor LDN-193189 (100 *nM*) and the GSK3 inhibitor CHIR-99021 (3 *µM*) from stage HH1 onwards (A-F) and with the VEGF receptor inhibitor Axitinib (100 *nM*) as described in methods. Layout of panels figure and scale bars as described in Fig. 1. Sections in C show the formation of extensive tissue folds after LDN-193189 and CHIR treatment.

Previous work has established how cells in the sickle-shaped posterior territory form the primitive streak. Initially cell-cell signalling between cells in the extra- and embryonic areas results in the induction of mesendoderm precursors in a sickle-shaped domain in the posterior embryonic epiblast (9). These cells then start a myosin II-dependent directional cell-cell intercalation process that drives the convergence and extension of the sickle-shaped mesendoderm in the extending primitive streak (6, 10, 11). This developmental phase is characterised by large counter-rotating vortical flows in the epiblast (Fig. S1B), known as polonaise movements (12, 13). During this process, the mesendodermal precursor cells begin to contract apically in preparation for their ingression in the streak (14, 15). Following an epithelial to mesenchymal transition (EMT) and ingression, the cells migrate away as cohorts of loosely associated cells to form the various endodermal and mesodermal structures in the embryo (Fig. 1C) (16).

These findings suggest that directed cell intercalation and cell ingression in the mesendoderm territory drive the morphogenetic flows in avian gastrulation. To test this idea, we inhibited mesoderm differentiation by blocking FGF signalling using the pan-FGF receptor inhibitor LY2874455. The addition of LY2874455 resulted in the complete inhibition of mesoderm differentiation and in the loss of the characteristic tissue flows associated with streak formation, (Fig. S2, movie 2). Analysis of the strain rate shows that there is little organised intercalation, confirming that the presence of mesendoderm tissue is a prerequisite for streak formation.

In contrast, the addition of an excess of FGF results in the circular expansion of the mesendoderm territory (9). During normal development SNAI2+ mesendoderm cells travel from the sickle-shaped territory where they are specified towards the primitive streak where they undergo EMT and cell ingression (14, 17). This can be seen via the presence of SNAI2+ cells in the inner layers and the breakdown of the fibronectin layer in the streak (Fig. 1B,C). Embryos treated with excess FGF show SNAI2+ cells have extended into an ingressed along a ring shaped region along the marginal zone (Fig.1 H,I,J), while the fibronectin basal layer is degraded. In-vivo imaging shows that FGF treated embryos exhibit a ring of tissue that presents the two characteristic signatures of the late primitive streak: cell ingression and perpendicular cell intercalation (Fig.1 K, L, M, movie 3). After ingression, the cells migrate towards the centre of the epiblast (movie 4). The resulting circular streak structure strongly resembles the germ band of teleost fish gastrulation, where cells undergo ingression and migrate towards the animal pole (18) (See also Fig. 3F and movie 9 of (8) for the corresponding DM.)

**Fig. 3:**
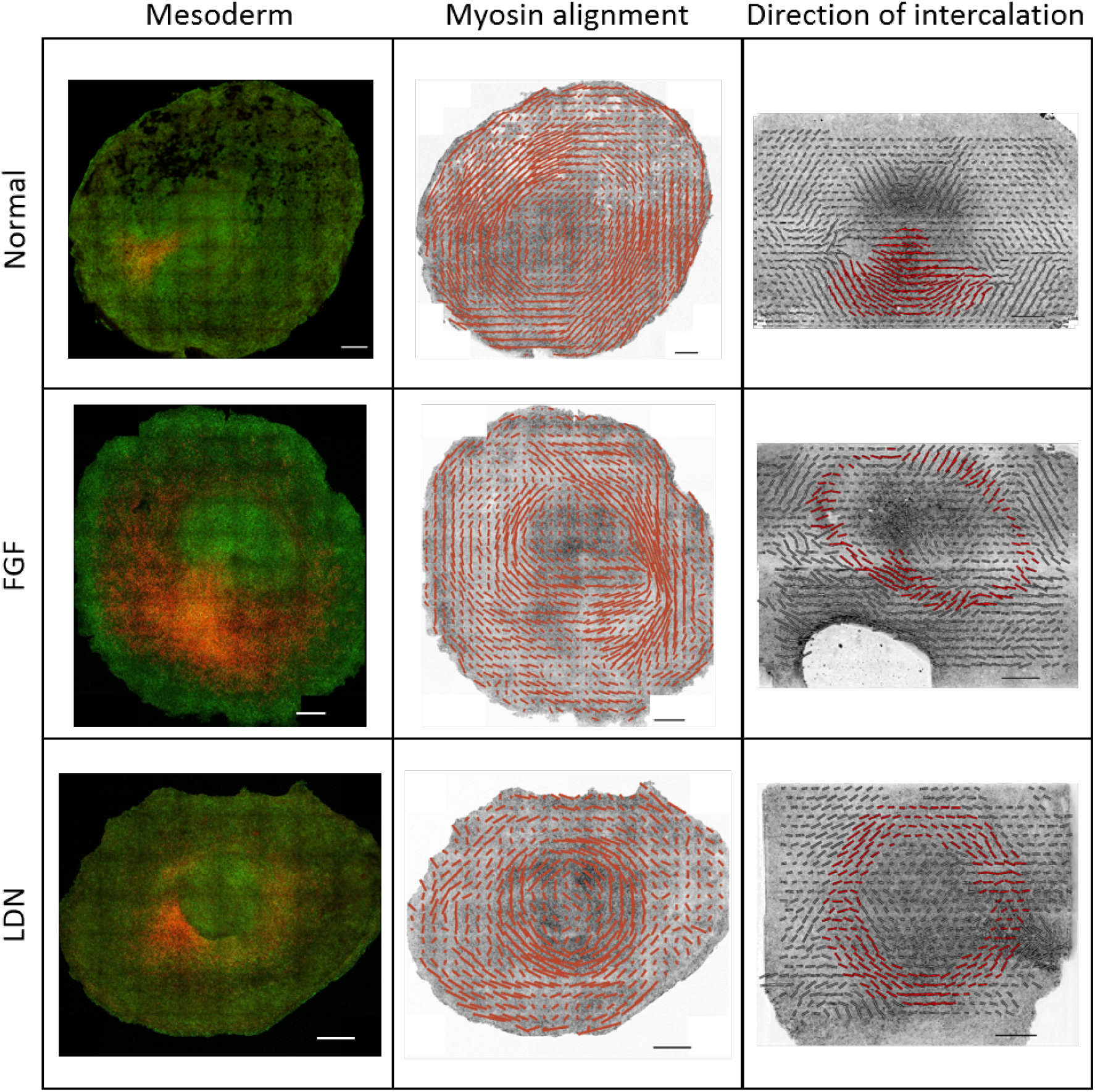
Comparison of patterns of mesendoderm differentiation and localisation (SNAI2+ cells in red), myosin alignment and direction of cell intercalation indicated by the anisotropic component of the P tensor (22). Top row: Control embryo after 16 hrs incubation in EC culture as described in methods. Second row: Embryo treated with FGF2 for 16hrs. Third row: embryo treated with LDN-193189 (100 *nM*) for 16hrs. Regions of intercalating mesendoderm cells are highlighted in red. For all images the scale bars represent 500um. For the myosin alignment (second column) the scale bars represent ∼ 30% anisotropy. For the direction of intercalation (third column) the scale bar represent a magnitude of ∼ 0.4 1*/h*.

Complementing the FGF-driven manipulation of the mesendoderm, we used a different molecular treatment based on BMP signalling inhibition in the anterior embryo (19) to reproduce the expansion of the mesendoderm territory. Application of the Alk2,3 receptor inhibitor LDN-193189 (Fig. S3A-D) allows us to de-repress the formation of mesoderm in the anterior region of the embryo, as seen by the expression of SNAI2+ cells in a circular domain (Fig. S3C). However blocking Nodal receptors Alk4,5 by the inhibitor SB50124 does not block mesoderm formation(Fig. S4). These mesoderm domains form rapidly and are more circular than after FGF addition, suggesting that relieving inhibition of mesoderm differentiation by LDN-193189 is faster than induction of mesendoderm by FGF. Concomittantly, we note the appearance of a circular fold at the boundary of the domain which contracted over time, engulfing the internal epiblast. The observed tissue buckling after blocking BMP signaling results from the inhibition of EMT and cell ingression, even in the presence of exogenous FGF (Fig. S3 G, H, I). Closer inspection of the development of these LDN-193189 embryos showed that they are noticeably smaller due to a significant inhibition of cell proliferation by BMP inhibition (Fig. S3E), preventing longer-term observation of the embryos in the light sheet microscope. To decouple the effects of ingression and proliferation we sought to restore cell proliferation using a combination of the BMP signalling inhibitor LDN-193189 and the GSK3 inhibitor CHIR-99021 (CHIR), conditions that have been used to produce mesendoderm precursors from IPS and ES cells (20). Treated embryos still showed the formation of mesodermal precursor rings and inhibition of EMT, resulting in buckling and involution of the central tissue (Fig. 2A-G), but without inhibition of cell division (Fig. S3F). Sectioning revealed that tissue in this internal domain had folded inside the embryo, and the mesendoderm cells internalised through the circular fold failed to undergo EMT but stayed connected in an epithelial sheet. Analysis of the anisotropic strain rate (Fig. 2E) in these embryos showed a pronounced contraction oriented tangential to the ring produced by cell intercalation (movie 5). This tangential intercalation drives ring contraction and a reduction in its circumference. Since the tissue inside the ring did not undergo EMT, it buckles producing the complete invagination of the epiblast as seen in cross-sections of embryos undergoing the same treatment (Fig. 2C). Interestingly, this situation mimics the tissue organisation and flow during the closing blastopore in amphibians, where tissue is invaginated as a continuous epithelial sheet (21) (see also Fig. 3H and movie 12 of (8) for the corresponding DM).

In addition to controlling the size and shape of the mesendoderm, we manipulated cell behavior by inhibiting VEGF signalling by using the inhibitor Axitinib (Fig. 2H-N) or competitive VEGFR2-Fc fragments (Fig. S5). When applied early, both treatments results in the formation of a short fat streak. Observations of the sickle-shaped mesendoderm showed that cell intercalation was not significantly inhibited, and the vortical flow of tissue into the ventral midline of the embryo continued. However there was a distinct absence of ingression of cells in the streak evidenced by the absence of tissue contraction in the midline (Fig. 2L, Fig. S5A,B) and lack of fibronectin degradation (Fig. S6), even though these manipulations do not effect cell divisions (Fig. S5C, D). Due to the lack of cell ingression, there are more cells flowing into the central streak area than flowing out of it. This results in the partial buckling of the epiblast at the anterior streak movie 6. There was no effect on cell divisions (Fig. S5C, D), suggesting that buckling was not the result of reduced division resulting in reduced tissue fluidity (23). The resulting fold structure resembles the blastoporal canal observed in reptilian gastrulation (24, 25) (see also Fig. 3D and movie 7 of (8) for the corresponding DM.)

Our experiments show that the the shape of the mesendoderm together with cell intercalation and ingression behaviours determine the morphogenetic outcome of gastrulation. While cell intercalations drive the main in-plane tissue flows, the presence or absence of individual cell ingression determine if the tissue will remain flat or will buckle. During normal development, the direction of intercalations of mesendoderm cells are strongly correlated with the direction of cables of phosphorylated myosin II that span several cells (6). At early gastrulation stages these myosin cables extend from posterior to anterior creating a semicircle between the embryonic and extraembryonic territories and they correlate with the direction of intercalation, as shown by the P tensor (22) computed from cell tracking data (Fig. 3) (6). We show that this strong correlation between mesendoderm expression domain and myosin planar cell polarity persists in the cases of circular mesendoderm domains, the FGF2 and LDN-193189 treatments (Fig. 3B,C). As in normal early development, the myosin planar cell polarity is initially oriented along the long axis of the mesendoderm expression domain, in the same direction of cell intercalations, which explains the contraction of the ring in the LDN + CHIR case. However, during normal streak extension at late stages (HH3-HH4) cell intercalations orient perpendicular to the long axis of the mesendoderm (primitive streak) driven by cell ingression in the streak. The same process also occurs in embryos treated with FGF, where many cells ingress through the circular streak, which results in the reorientation of the direction of intercalation of nearby cells perpendicular to the ring. These observations show that FGF induces mesoderm that will show both intercalation and ingression, while LDN-CHIR results in relief of mesoderm inhibition as well as an inhibition of EMT.

To quantify these observations, in the accompanying paper (8) we propose and explore a theoretical model for gastrulation as a mechanosensitive self-organising process. Our continuum model couples the myosin driven tension-sensitive active stress to tissue flow and cable ordering, and leads to an interpretation of gastrulation as the result of a spontaneous instability in the early embryo. Our model can recapitulate these different gastrulation modes by varying the extension of the mesendoderm and a parameter modulating the amount of active cell ingression (see Fig. 2-4 and movies 2-12 of (8)), showing that a tension-dependent self-organisation of the tissue flows is sufficient to drive the different gastrulation morphogenetic programs.

Surprisingly, despite being an essential phase of early development, the exact morphology of gastrulation shows considerably variation across the evolutionary tree. Furthermore, dissociated and re-aggregated embryonic stem cells (ESCs) can re-establish a body axis despite the loss of the pre-existing positional information, showing that gastrulation is a highly regulative self-organising process (27). Here we have shown that directed cell intercalations and cell ingressions within the mesendoderm generate the tissue flows driving avian gastrulation. Furthermore by targeting key developmental signalling pathways, we uncoupled cell intercalation and ingression and modulated the size of the mesendoderm territory. These manipulations are sufficient to induce large morphogenetic changes that recapitulate the hallmark morphologies of different vertebrate gastrulation modes in a single organism, the chick embryo. These results suggest that the modulation of the extension of the mesendoderm territory and the presence or absence of EMT controlling cell ingression largely determined the different morphologies observed during the evolution of vertebrate gastrulation (Fig. 4). Altogether, these results suggest that different gastrulation modes could evolve because early development is a flexible self-organising process, where quantitative changes in a relatively small number of signalling pathways can drive major morphogenetic changes though their effects on modulating individual cell behaviours.

**Fig. 4:**
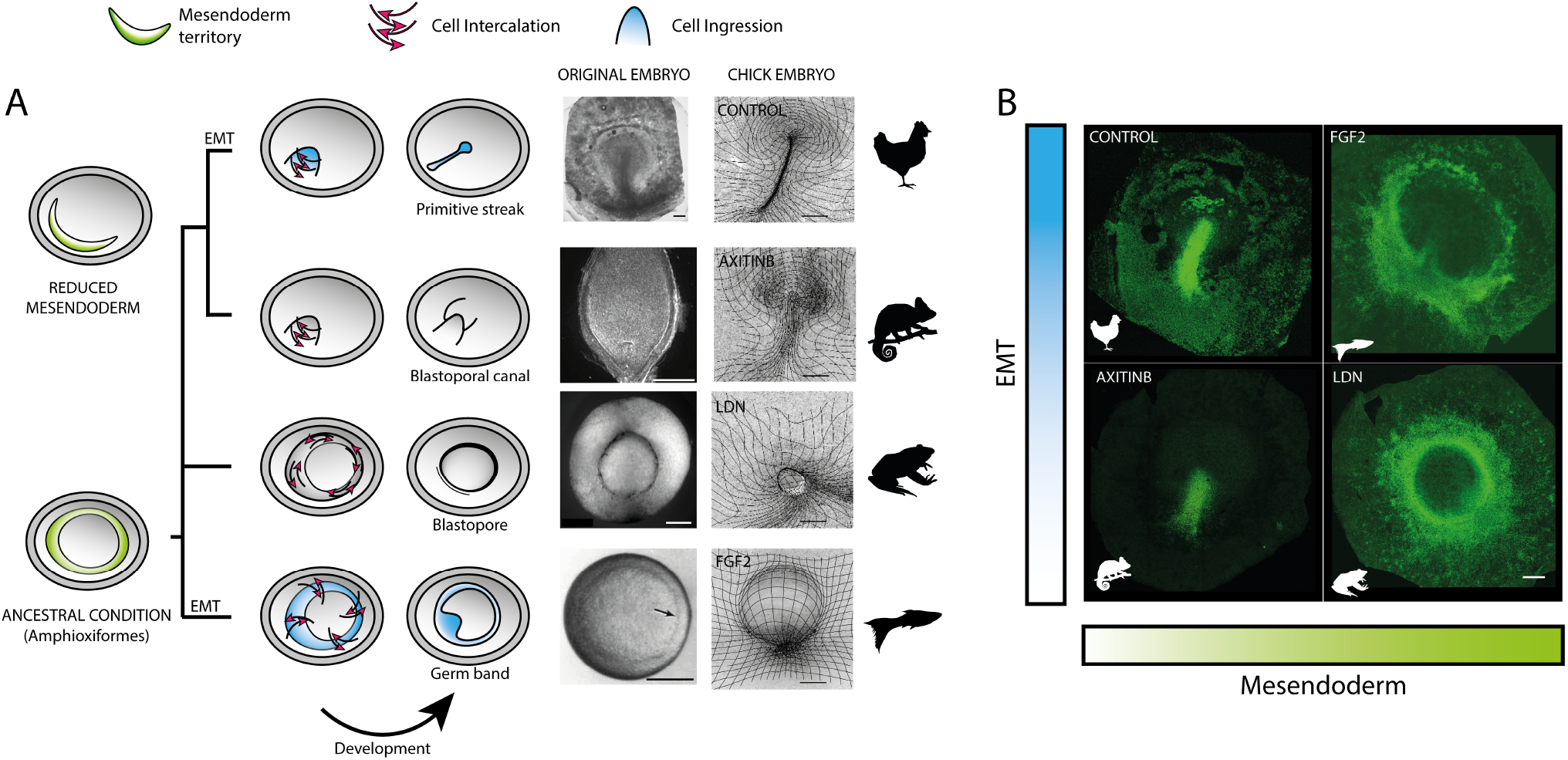
Changes in the extent of the mesendoderm territory and the degree of EMT could underlie the evolution of vertebrate gastrulation. A: Evolutionary relationships of representative vertebrate morphologies recapitulated in the chick embryo via modification of key cell behaviours. Images of gastrulation in reptiles (d28), amphibians (12h) and fish (5.7h) adapted from (21, 24, 26), scale bars 250 *µm*. B: SNAI2+ cell distributions in HH3+ chick embryos after different treatments show that manipulating the extent of the mesendoderm territory and the level of EMT controlling ingression in the chick embryo reproduce critical morphological structures of gastrulation in other vertebrates. Scale bars for chick embryos 500 *µm*.

## Acknowledgments

GSN acknowledges support from an EASTBIO BBSRC PhD student training grant (1785593). MS acknowledges the Schmidt Science Fellowship and the Postdoc Mobility Fellowship from the Swiss National Foundation for partial support. LM thanks the NSF-Simons Center for Mathematical and Statistical Analysis of Biology Award 1764269, NIH 1R01HD097068, the Simons Foundation, and the Henri Seydoux Fund for partial support. CJW thanks the BBSRC (BB/N009789/1, BB/K00204X/1, BB/R000441/1, BB/T006781/1) for financial support and a Wellcome Trust imaging equipment award (101468/Z/13/Z) for partial support.

## Supplementary Information for

### Material and Methods

#### Chemicals and other reagents

LDN-193189, CHIR-99021, Axitinib, SB50124 were acquired from Sigma. Hydromount was obtained from National Diagnostics. Ultrapure grade was acquired from Merck/VWR. Recombinant Human FGF2 (233-FB) and Recombinant Human VEGFR2/KDR-Fc (357-KD-050) were obtained from (R&D Systems). Primary antibodies: Phospho-myosin light chain 2 (S19) mouse mAB (3675), Phosphomyosin light chain 2 (T18, S19) rabbit mAB (3674), Slug(C19G7) rabbit mAB(9585) were from Cell Signalling Technology. Alexa Fluorophore conjugated secondary antibodies were obtained from Invitrogen.

#### Chick lines and Embryo culture

Fertile eggs from White Leghorn chicks were obtained from Henri Stewart&Co Lincolnshire UK. Membrane GFP embryos were obtained for the National Avian Research facility Edinburgh University. Embryos were cultured in EC culture according to published procedures (1).

#### Imaging

To image the complete the chick embryo during gastrulation we use a chick line expressing GFP in the cell membranes (myr-GFP) and dedicated light-sheet microscope (2). To cover the entire embryo (∼ 4 mm in diameter), we acquire images at intervals of 1.84 *µm* at 45° with respect to the surface in 2 overlapping sequential scans (∼ 5000 images of 2560 × 400 pixels) covering a roughly a 3.2 mm x 4.7 mm area every 3 minutes. Embryos are cultured during imaging as described in (3). In these conditions, chick embryos can develop for more than 24 hours at 37°C (480 time-points, *>* 2TB).

#### Image processing

##### Surface focusing

First, we transform the image volumes from the original 45° geometry to a 90° geometry. Once the image volumes have been transformed, we find the surface using a method based on the square gradient focusing algorithm (4). Briefly, we tile the volume in 25×25 pixel columns and use the algorithm to find the position of the surface in each column, which constitutes the height map. Finally we apply a smoothing filter to the height map, and use this to section the image volume to produce a 2D image of the surface of the embryo (Fig. S 1 A).

##### Velocity fields and strain rates

The velocity fields at the surface of the embryo is computed by digital particle image velocimetry (PIV; Fig. S 1 B) using PIVLab vs1.32 for MATLAB (5) with two passes of 64 × 64 pixels and 32 by 32 pixels with a 50% overlap as in (6). The gradient of the velocity can be used to compute the strain rate at the surface of the embryo. They can be decomposed into two parts: the isotropic term which describes the local rate of change in the area, and the anisotropic term which describes the local rate shear deformation. Together they can be used as an indirect, but robust measure of cell behaviours. In a confluent epithelial sheet, as the epiblast, the isotropic term is the product of the balance of changes in the cell area, cell divisions and cell ingressions. In contrast, the anisotropic part is the result of directed cell intercalation or coordinated asymmetric changes in cell shape (Fig. S 1 C). The implementation details can be found in (6).

##### Division detection

We developed a new robust algorithm to detect and track individual cell divisions independently of image segmentation. During cytokinesis, epithelial cells round up on the surface, acquiring a very typical circular shape. We used the Circular Hough Transform (CHT) implementation in MATLAB to find the characteristic circular shapes associated with dividing cells. The algorithm uses PIV to follow the dividing cells to avoid counting the same division event repeatedly between consecutive frames.

#### Myosin quantification

To quantify the myosin concentration and alignment the surface was identified using the above described procedure, using typically the actin signal to finding the apical surface (Fig. S 1 E, F). This surface projection was analysed for myosin anisotropy using a Fourier transform based algorithm (Fig. S 1 G) to find cellular anisotropy and size using a sub-tiling of the surface in 256*256 pixels and a 50% overlap between tiles (7).

#### Chemical manipulations

For in-vivo imaging, the embryos were later transferred to imaging chambers as described in (3), and the chemical inhibitors were dissolved in the 10 mL albumen that cover the embryos during imaging. For immunocytochemistry experiments chemical inhibitors were dissolved in the EC culture substrate. Recombinant Human FGF2 and VEGFR2-Fc were added as a 1 *µL* drop (50 *µg/mL*) carefully deposited on the hypoblast side of embryos in EC culture. For life-imaging, the embryos were later transferred to imaging chambers maintaining the added protein on the hypoblast side of the embryo.

#### Immunocytochemistry

After the chemical manipulations, the embryos were fixed in 4% paraformaldehyde/PBST (PBS containing 0.1% Tween20) on ice during 3h and later washed them three times in PBS. Later the antibody detection was performed as described (8) followed by staining with DAPI and Alexa 405 Phalloidin. Embryos were mounted in Hydromount and image using a wide field microscope and or a Leica TCS SP8 confocal microscope at 10-20x magnifications. Selected embryos were processed for sectioning, first embryos were embedded in 7.5% gelatine/15% sucrose, followed by freezing in a dry ice/isopentane mixture and sectioned on a Leica cryostat. Sections were mounted on lysine coated microscope slides and mounted with Hydromount.

#### Confocal microscopy

To image the embryos confocal tile scans were taken at 10x and 20x magnification using sequential line scanning on a Leica SP8 confocal microscope. To cover the thickness of the embryo stacks of 20-40 slices were taken at 2-4um intervals along the z-axis

#### Dynamic Morphoskeletons

Given a planar velocity field v(x, *t*), we compute the Dynamic Morphoskeleton (DM) (9) from the backward and forward Finite Time Lyapunov Exponents (FTLE). We compute the FTLE as

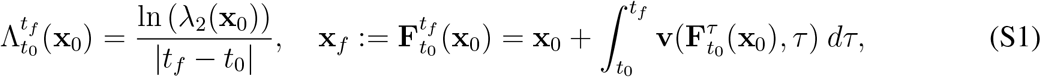

where *λ*_*2*_ (x_0_) denotes the highest singular value of the Jacobian of the flow map 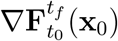 and 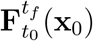 the flow map describing the trajectories from their initial **x**_0_ to final **x**_*f*_ positions. To compute the FTLE, we first calculate 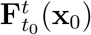 by integrating the cell velocity field **v**(**x**, *t*) using the MATLAB built-in Runge-Kutta solver ODE45 with absolute and relative tolerance of 10^*−*6^, linear interpolation in space and time, and a uniform dense grid of initial conditions. Then, denoting the *i* − *th* component of the flow map 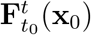 by 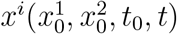, we compute the deformation gradient 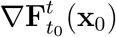 using the finite-difference approximation

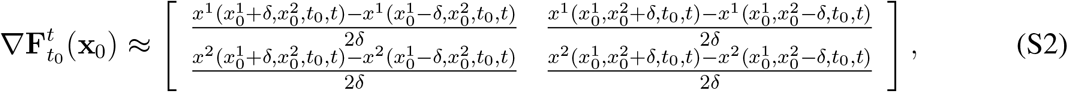

where *δ* is the initial conditions’ grid spacing. After computing 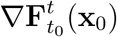, we use eq. (S1) for computing the FTLE field.

## Supplementary figures

**Fig. S 1:**
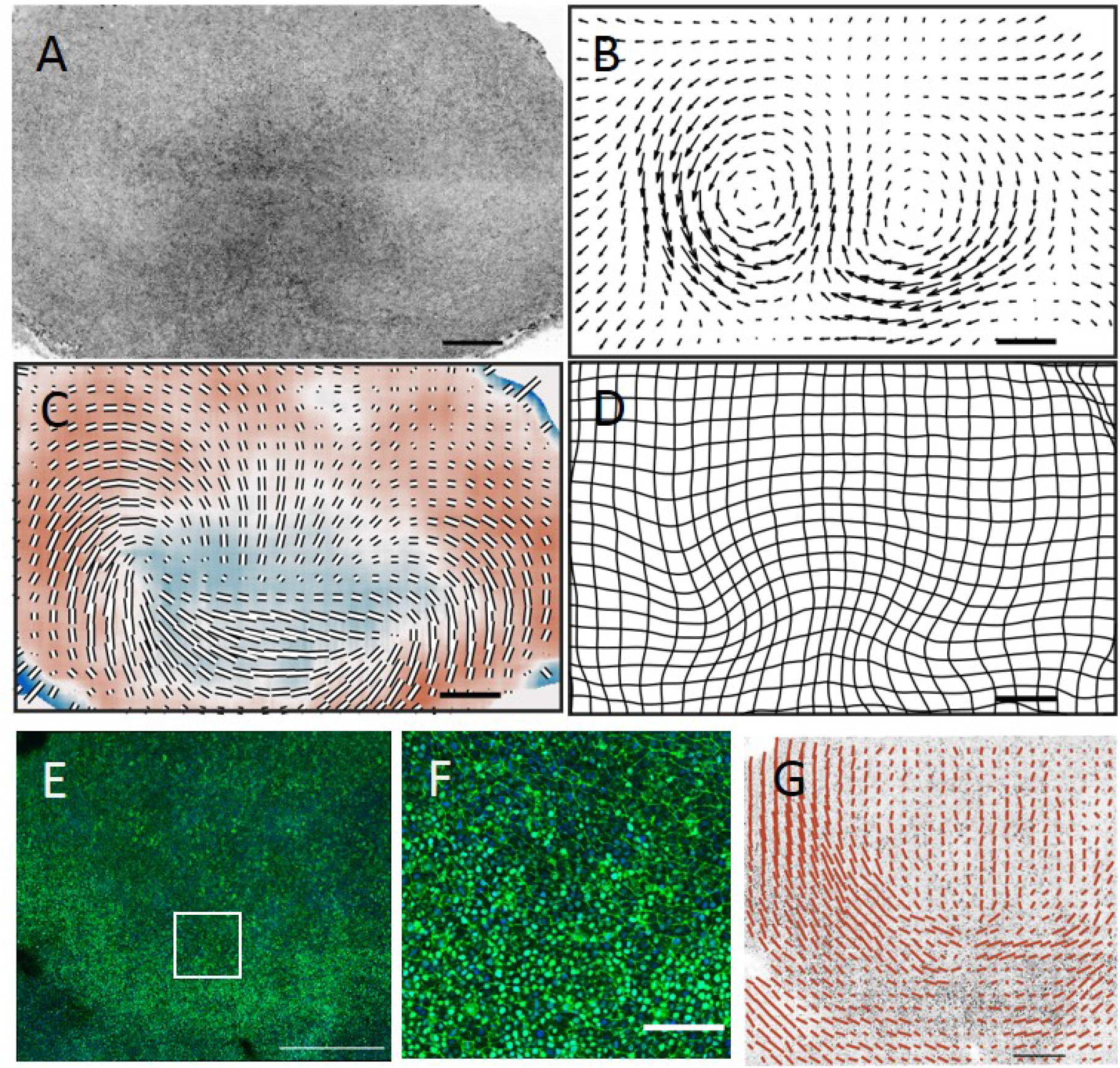
Correlation of tissue flow, deformation speed, cell shape and myosin distribution and alignment. A: Epiblast view of HH2 stage embryo in lightsheet microscope B: velocity field for embryo (HH2) shown in A. C: Strain rate tensor, isotropic component colour coded red blue, anisotropic component shown as black bars in drection of contraction. D: Deformation grid calculated from stage HH1 to current stage HH2 for embryo shown in A. D: Slug and phosphomyosin light chain in HH2 stage embryo. E: High magnification image of section in white box shown in D showing myosin cables and nuclear slug staining. Myosin anistropy computed for embryo shown in D. Scale bars in A-F 500 *µm*. In C the scale bar length represents ∼ 0.5 1/h for the anisotropic component of the strain rate. In G the scale bar represents 250 *µm* and ∼ 70% anisotropy

**Fig. S 2:**
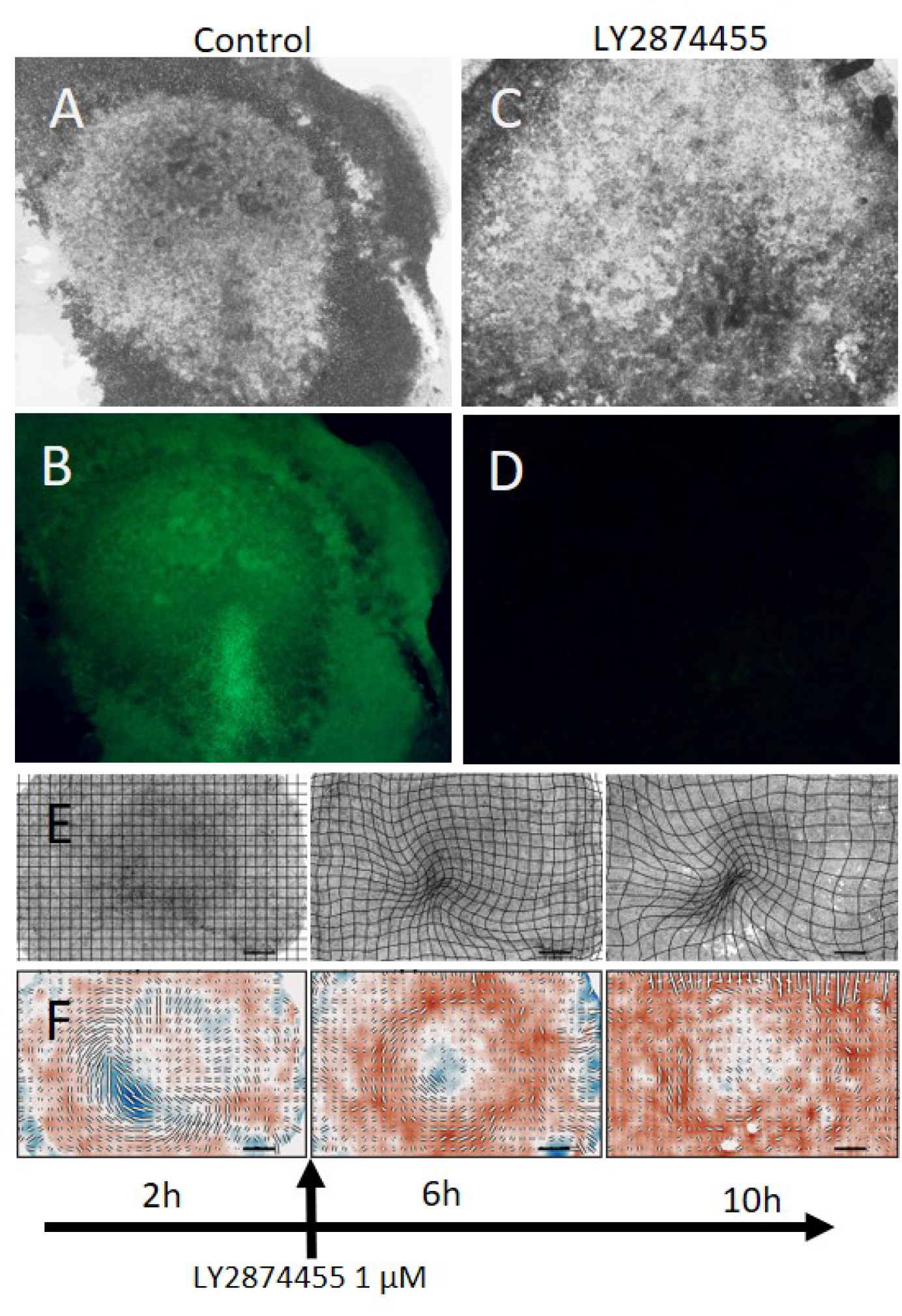
Inhibition of FGF signalling blocks mesoderm differentiation and streak formation. A: surface view of control embryo after 16hrs of development in EC culture. B: Slug expression of embryo shown in A. C: Embryo after 16hrs of treatment with the FGF receptor inhibitor LY2874455 (1uM). D: absence of Slug expression in embryo shown in C. E: Three successive time of embryo images in lightsheet 4, 8 and 12 hours faster addition of LY2874455 (1uM). Note the absence of the formation of a clear streak. F: strain rate tensor at 4,8 and 12 hours for embryo shown in C. Scale bars for E and F as in Fig. S1 C

**Fig. S 3:**
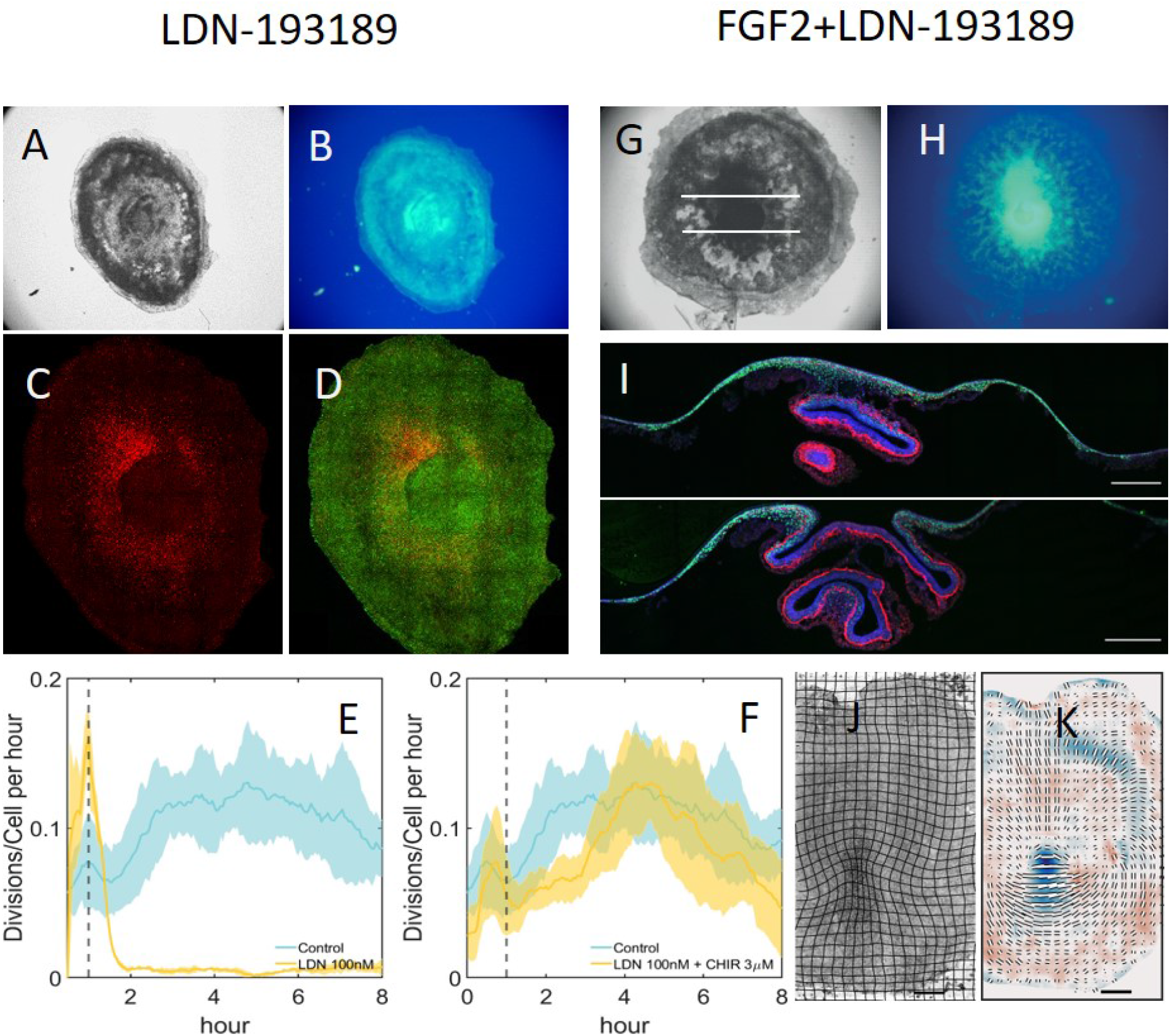
Changes in gastrulation morphology after interfering with BMP and FGF signalling. A: widefield image of embryo treated for 16hrs with LDN -193189 for 16hrs. B: Widefield image of (S19)phospho-myosin light chain expression of embryo shown in A. C: Confocal image of slug expression of embryo shown in A. D: Confocal image of (S19) phospho-myosin light chain and slug expression in embryo shown in A. E: Cell division rate in embryo during LDN -193189 treatment (100nM) measured in lightsheet microscope compared to control (both curves shown means and standard errors of division rates in three embryos). F: Cell division rate in embryo during LDN-193189 plus CHIR (3 *µM*) treatment measured in lightsheet microscope, compared to control (both curves shown means and standard errors of division rates in three embryos). G: widefield image of embryo after treatment of LDN-193189 and FGF2 for 16hrs. I: Widefield image of (S19) phospho-myosin light chain expression of embryo shown in G. I: Two sections through embryo shown in G at position of white lines. J: lightsheet microscopy images of embryo treated with LDN -193189 for 10 hours. K: Strain rate tensor for embryo shown in J. Scale bars as in figure 1, except in I where the scale bars are 250 *µm*. Scale bar for K as in Fig. S1 C

**Fig. S 4:**
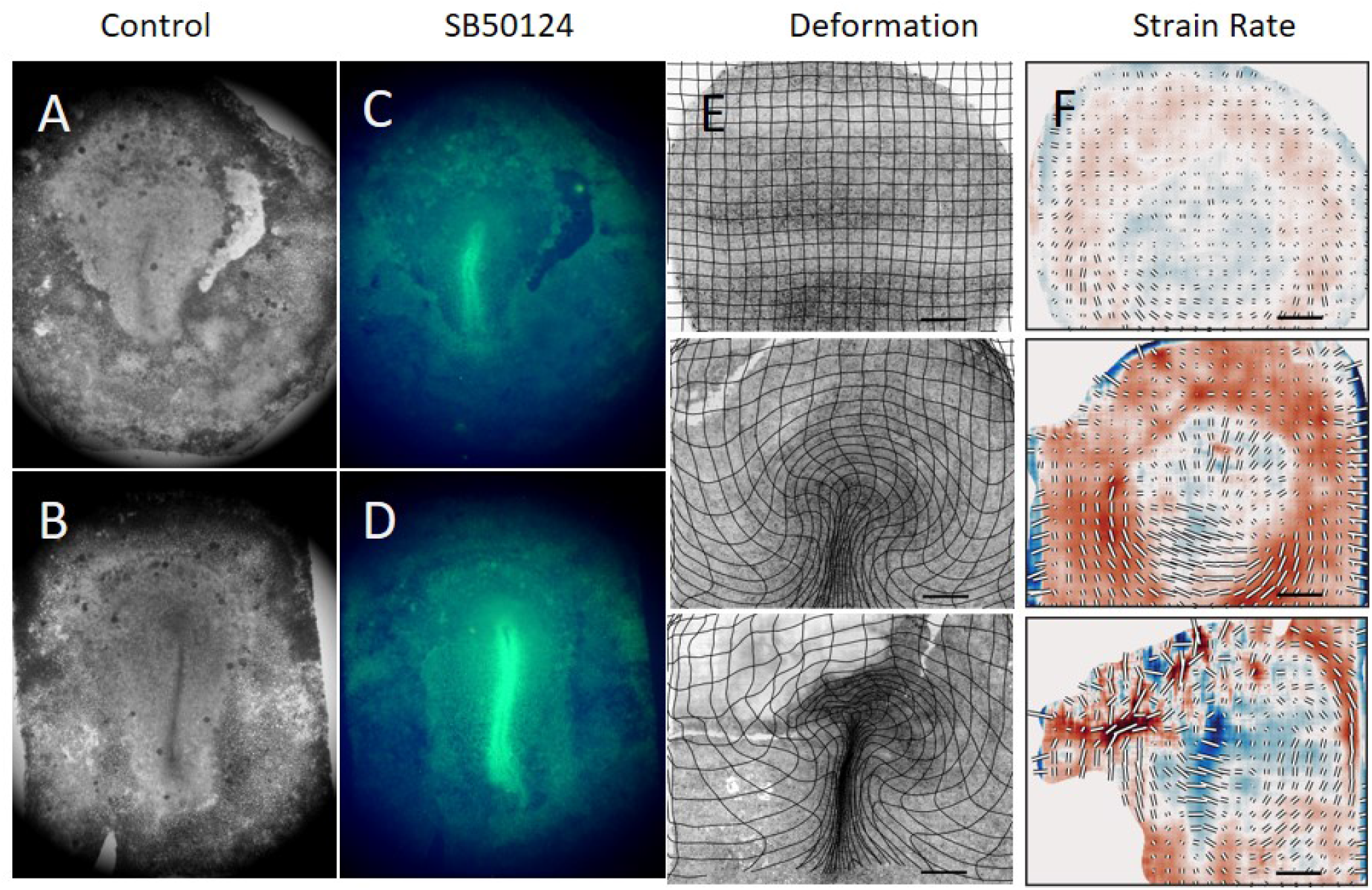
Inhibition of Nodal signalling partly inhibits streak formation. A: widefield image of control embryo after 16 hrs development in EC culture. B: Slug expression in embryo shown in A. C: Embryo after 16 hrs incubation in EC culture with the Nodal signalling inhibitor SB50124 (10 *µM*). E: Surface view of embryo in light sheet microscope 2, 4 and 8 hours the after addition of SB50124 (10 *µM*). F: Strain rate tensor of the embryo at successive times shown in E. Scale bars 500 *µm*. Scale bars for F as in Fig. S1 C

**Fig. S 5:**
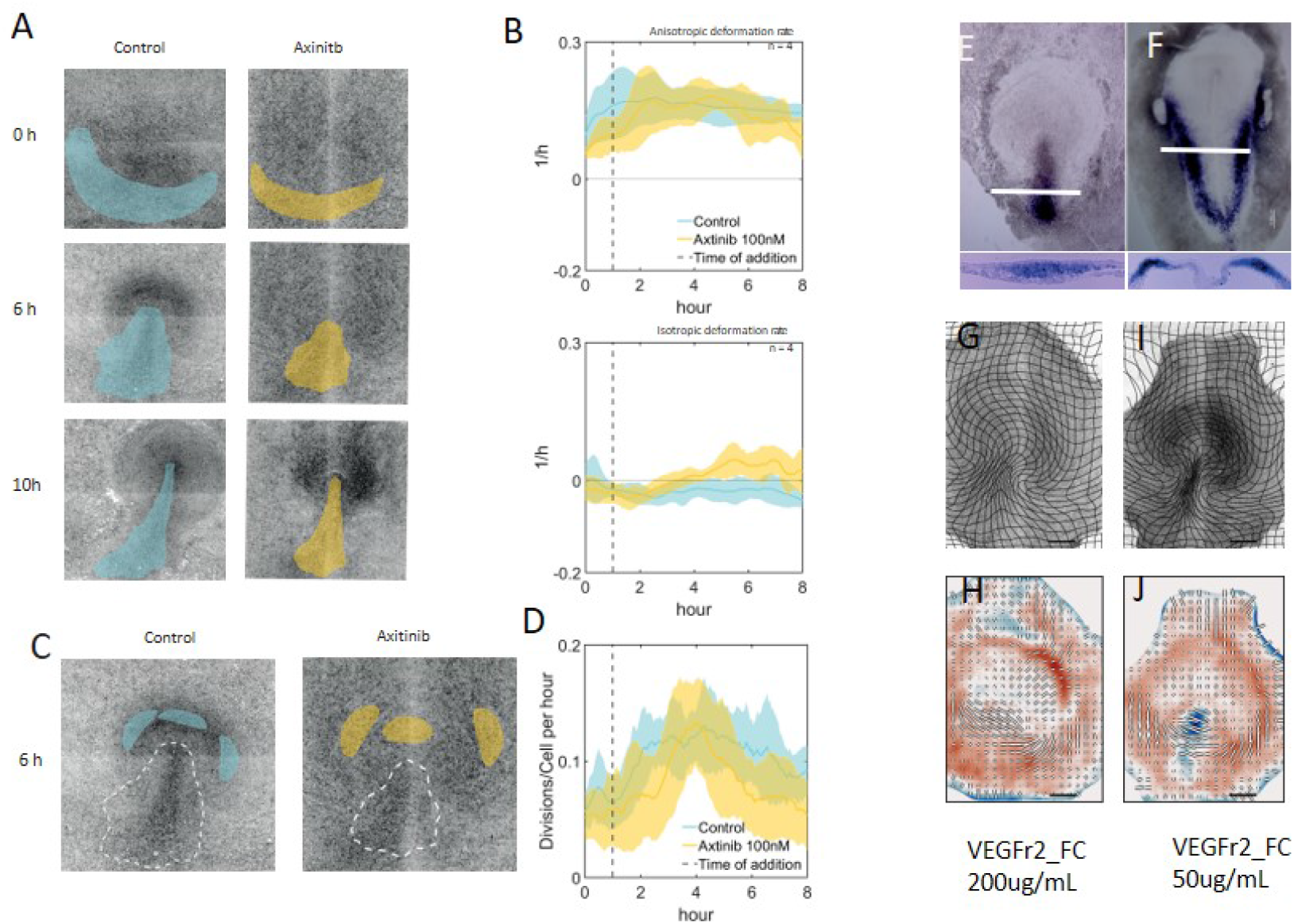
Inhibition of VEGF signalling results in loss of ingression but does not inhibit intercalation. A: Comparison of control embryo Left column and an embryo treated with the VEGF signalling inhibitor Axitinib (100nM) (right column). Images are overlaid with areas that correspond to the mesendoderm territories as determined from the dynamic morphoskeleton for the control embryo (blue) and axitinib embryo (yellow). B: Strain rate tensor quantified in the blue and yellow areas in A, showing that in the control case the isotropic strain rate (upper panel) becomes negative, while in the axitinib treated embryos the isotropic component stays positive shown a lack of contraction, while the anisotropic strain component is high in both cases showing significant intercalation. The curves shown represent the mean and standard error of three embryos in each case. C,D: Analysis of cell division rate in selected areas of the anterior epiblast of control and axitinib treated embryos. For each embryo three analysis regions anterior to the streak (C) were followed over time and the rate of cell division (D) calculated. The results show that axitinib treatment does not affect the rate of cell divisions. The curves in D are the average and standard errors of three regions for three embryos for each treatment. E, In situ hybridisation of the VegfR2 receptor at two stages of development (HH3 and HH5), sections show that the receptor is expressed in mesendoderm before and after ingression. G: Epiblast view and defromation grid of embryo treated for 16hrs with VegfR2 Fc fragments to deplete its ligands in the ligthsheet microscope. H: Strain rate tensor for embryo shown in H. Note the presence of intercalation of a strong anisotropic component (black bars) showing intercalation, but an absence of a negative isotropic component (blue), showing a lack of ingression. I,J: Lower VegfR2-Fc concentrations show a partial inhibition of streak formation. Scale bars for H, J as in Fig. S1 C

**Fig. S 6:**
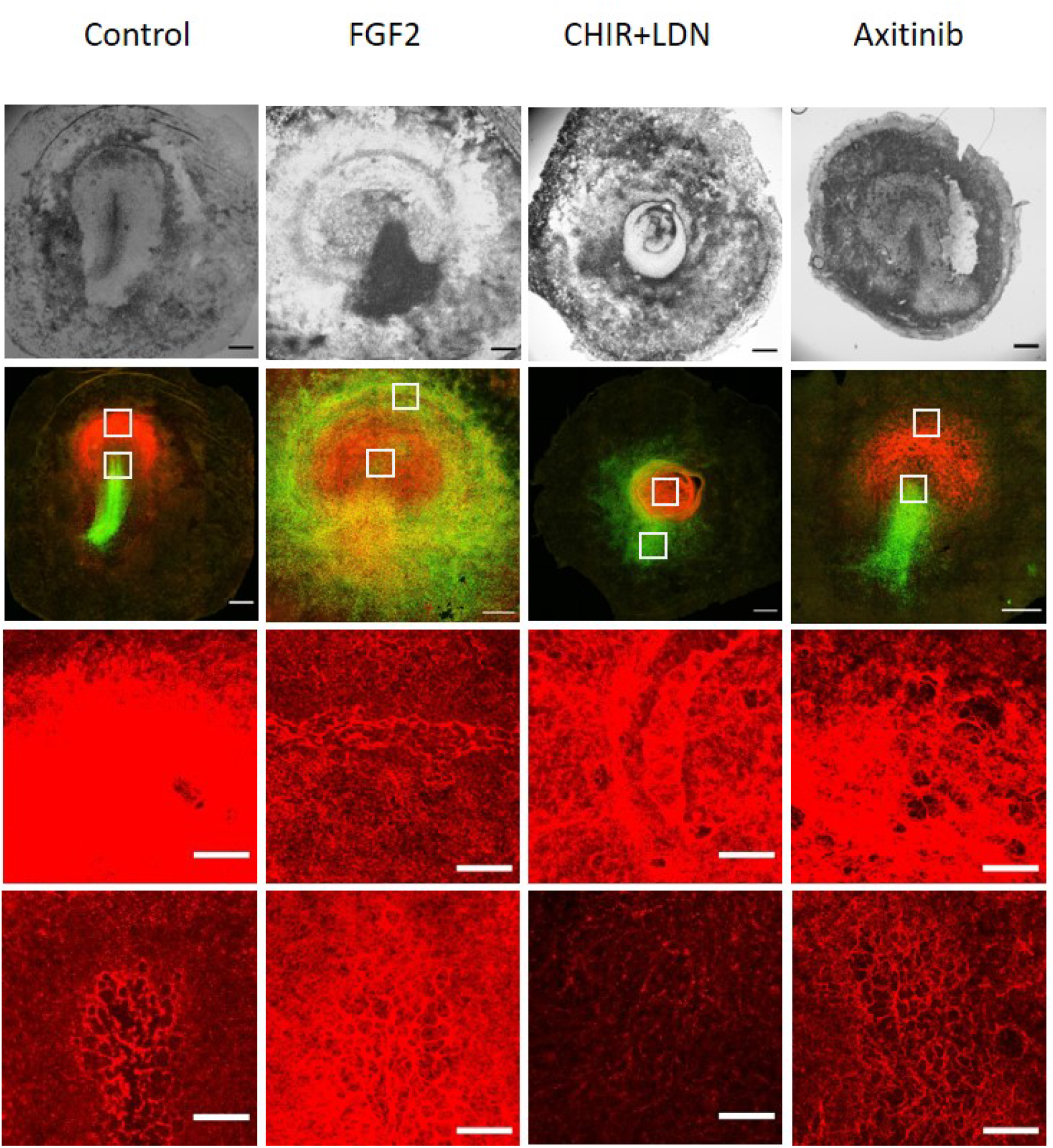
Comparison of slug and fibronectin expression after various perturbations. Top row: Brightfield images of embryos after 16hrs in EC culture after various treatments. Second row: Confocal images of the embryos in the top row showing expression of Slug (green) and fibronectin (red). The third and fourth rows show higher magnification images takes in front of the streak (upper white square in each image) and (lower white square in each image) at the tip of the streak. It can be seen in the control (first row) that the region anterior to the streak expresses high levels of fibronectin forming a densely packed meshwork of fibrils. In the streak the fibronectin shows a larger pore meshwork through which the cells ingress. Note that in the ring shaped streak after FGF treatment (second column) the fibronectin shows the same large porous structure as in the streak, indicating that this also a region where the cells ingress. After LDN and axitinib treatment the anterior region shows a dense fibrillar network while in the streak or ring cases the porous network is less well developed as in the control, or after FGF treatment. The data show th1a1t the presence or absence of ingression is reflected in the fibronectin network structure. Scale bars in two top rows of images 500 *µm*, scale bars in two bottom rows of images 100 *µm*

## Supplementary movies

Red titles are hyperlinks to the movies. Click to open the movies in the browser.

Movie 1. Development of control embryo from stage HH1 to HH3+. Movie shows bright field image (top panel) and strain rate tensor of the same embryo (bottom panel) Isotropic strain rate coloured blue (contraction) to red (expansion) scale bar 500 *µm*. Time interval is 3 minutes.

Movie 2. Development of embryo treated with FGF signaling inhibitor LY287455. Movie shows bright field image (top panel) and strain rate tensor of the same embryo (bottom panel) Isotropic strain rate coloured blue (contraction) to red (expansion) scale bar 500 *µm*.

Movie 3. Development of embryo treated with FGF2. Movie shows bright field image (top panel) and strain rate tensor of the same embryo (bottom panel) Isotropic strain rate coloured blue (contraction) to red (expansion) scale bar 500 *µm*.

Movie 4. Migration of mesoderm cells after addition of FGF2. Sections were taken deeper in the embryo show the mesoderm cells ingressing through the circular primitive streak are seen to be migrating towards the centre of the embryo.

Movie 5. Combined effect of CHIR+LDN on mesoderm induction and streak formation Movie shows bright field image (top panel) and strain rate tensor of the same embryo (bottom panel) Isotropic strain rate coloured blue (contraction) to red (expansion) scale bar 500 *µm*. Time interval is 3 minutes.

Movie 6. Effect of inhibition of Vegf signaling with Axitinib on streak formation Movie shows bright field image (left panel) and strain rate tensor of the same embryo (right panel) Isotropic strain rate coloured blue (contraction) to red (expansion) scale bar 500 *µm*. Time interval is 3 minutes.

Movie 7. Comparison of tip of streak formation in control embryo and embryo treated with 100 nM Axitinib. Time interval is 3 minutes.

## Notes

### Competing Interest Statement

The authors have declared no competing interest.

## References

1. L. Solnica-Krezel, Gastrulation: From embryonic pattern to form (Academic Press, 2020).

2. D. Arendt, K. Nübler-Jung, Mechanisms of development 81, 3 (1999).

3. D. Kimelman, Nature Reviews Genetics 7, 360 (2006).

4. T. Merle, E. Farge, Current opinion in cell biology 55, 111 (2018).

5. E. Hannezo, C.-P. Heisenberg, Cell 178, 12 (2019).

6. E. Rozbicki, et al., Nature cell biology 17, 397 (2015).

7. M. Serra, S. Streichan, M. Chuai, C. J. Weijer, L. Mahadevan, Proceedings of the National Academy of Sciences 117, 11444 (2020).

8. M. Serra, et al., bioRxiv (2021).

9. C. Alev, Y. Wu, Y. Nakaya, G. Sheng, Development 140, 2691 (2013).

10. O. Voiculescu, F. Bertocchini, L. Wolpert, R. E. Keller, C. D. Stern, Nature 449, 1049 (2007).

11. M. Saadaoui, D. Rocancourt, J. Roussel, F. Corson, J. Gros, Science 367, 453 (2020).

12. L. Gräper, Wilhelm Roux’Archiv für Entwicklungsmechanik der Organismen 116, 382 (1929).

13. M. Chuai, et al., Developmental biology 296, 137 (2006).

14. Y. Nakaya, E. W. Sukowati, Y. Wu, G. Sheng, Nature cell biology 10, 765 (2008).

15. O. Voiculescu, L. Bodenstein, I.-J. Lau, C. D. Stern, Elife 3, e01817 (2014).

16. X. Yang, D. Dormann, A. E. Münsterberg, C. J. Weijer, Developmental cell 3, 425 (2002).

17. H. Acloque, et al., Developmental cell 21, 546 (2011).

18. D. Pinheiro, C.-P. Heisenberg, Current topics in developmental biology 136, 343 (2020).

19. C. F. Arias, M. A. Herrero, C. D. Stern, F. Bertocchini, Scientific reports 7, 1 (2017).

20. J. Chal, et al., Nature protocols 11, 1833 (2016).

21. D. R. Shook, E. M. Kasprowicz, L. A. Davidson, R. Keller, Elife 7, e26944 (2018).

22. F. Graner, B. Dollet, C. Raufaste, P. Marmottant, The European Physical Journal E 25, 349 (2008).

23. J. Firmino, D. Rocancourt, M. Saadaoui, C. Moreau, J. Gros, Developmental cell 36, 249 (2016).

24. M. J. Stower, et al., Developmental Dynamics 244, 1144 (2015).

25. M. J. Stower, F. Bertocchini, Wiley Interdisciplinary Reviews: Developmental Biology 6, e262 (2017).

26. C. B. Kimmel, W. W. Ballard, S. R. Kimmel, B. Ullmann, T. F. Schilling, Developmental dynamics 203, 253 (1995).

27. N. Moris, A. M. Arias, B. Steventon, Current opinion in genetics & development 64, 78 (2020).

## References

1. S. C. Chapman, J. Collignon, G. C. Schoenwolf, A. Lumsden, Developmental dynamics: an official publication of the American Association of Anatomists 220, 284 (2001).

2. E. Rozbicki, et al., Nature cell biology 17, 397 (2015).

3. E. Rozbicki, W. Cj, Nature Protocol Exchange (2015).

4. A. M. Eskicioglu, P. S. Fisher, IEEE Transactions on communications 43, 2959 (1995).

5. W. Thielicke, E. Stamhuis, Journal of open research software 2 (2014).

6. A. I. Karjalainen, Automatic cell tracking for studying large epithelial sheets of cells and its application to chicken gastrulation, Ph.D. thesis, University of Dundee (2015).

7. M. Durande, et al., Physical Review E 99, 062401 (2019).

8. N. R. Leslie, X. Yang, C. P. Downes, C. J. Weijer, Current Biology 17, 115 (2007).

9. M. Serra, S. Streichan, M. Chuai, C. J. Weijer, L. Mahadevan, Proceedings of the National Academy of Sciences 117, 11444 (2020).

